# Deep Convolutional Network for Animal Sound Classification and Source Attribution using Dual Audio Recordings

**DOI:** 10.1101/437004

**Authors:** Tuomas Oikarinen, Karthik Srinivasan, Olivia Meisner, Julia B. Hyman, Shivangi Parmar, Robert Desimone, Rogier Landman, Guoping Feng

## Abstract

We introduce an end-to-end feedforward convolutional neural network that is able to reliably classify the source and type of animal calls in a noisy environment using two streams of audio data after being trained on a dataset of modest size and imperfect labels. The data consists of audio recordings from captive marmoset monkeys housed in pairs, with several other cages nearby. Our network can classify both the call type and which animal made it with a single pass through a single network using raw spectrogram images as input. The network vastly increases data analysis capacity for researchers interested in studying marmoset vocalizations, and allows data collection in the home cage, in group housed animals.

## I. INTRODUCTION

Convolutional neural networks (in 2D) have seen a lot of success in the fields of environmental and animal sound classification (Boddapati, Petef, Rasmusson, & Lundberg, 2017). Using convolution in both the time and frequency domain makes sense for these tasks since animal calls and environmental sounds often have distinct structure that can be clearly seen in spectrograms or other image representations of audio data. Here we present a neural network for auto detection, classification and attribution of vocalizations in the common marmoset (Callithrix Jacchus).

The impetus for this work is research using marmosets as a primate model to study mental disorders affecting social behavior, such as autism ((Jennings et al., 2016; Miller et al., 2016). In marmosets, vocal exchanges are an essential part of social interaction (Eliades & Miller, 2017), and are at least partly learned from parents and peers. Analysis of vocalizations can yield measures of vocal development, and vocal interactions can potentially be used to track sociability and learning of social rules. However, labeling vocalizations in audio recordings is labor-intensive, therefore automation is valued highly.

Our dataset consists of dual channel recordings from normal, captive marmoset monkeys housed in pairs, where each animal wears a voice recorder. The data is annotated by researchers for a variety of call types. We present a neural network that can detect and classify the calls from each animal with high accuracy based on the spectrogram. We use one network with two convolutional streams, which are concatenated in the end and followed by a single fully connected layer with dropout before a final softmax layer that classifies both the type and the source of the animal call.

In this paper, Section 2 covers background and previous models, Section 3 describes the details of the experiment, the dataset and the neural network model, in Section 4 we report the results, in Section 5 we present a discussion and conclusions.

## II. BACKGROUND AND RELATED WORK

The common marmoset (Callithrix Jacchus) is among the smallest primates and is gaining interest as a non-human primate model for neuroscience research (Jennings et al., 2016). The species lends itself well for studying social behavior, since marmosets have features in common with humans that are not found in every primate species, such as vocal interaction, imitation and cooperative breeding (Miller et al., 2016). In the vocal domain, marmosets have a repertoire of at least 8 call types, which occur in different conditions, and are thought to convey different information to others. For example, there are calls that serve to maintain contact with members of their group, calls that broadcast the presence of a threat and calls that signal inter-group threats (Bezerra & Souto, 2008; Miller, Mandel, & Wang, 2010). In both humans and marmosets, vocal interactions are structured and organized according to set principles (Sacks, Schegloff & Jefferson, 1974). The exchange of contact calls between marmosets shows a turn-taking dynamic that is comparable to turn-taking in human conversation and other interactions (Henry, Craig, Lemasson, & Hausberger, 2015; Levinson & Torreira, 2015).

We distinguish three functions that an automatic vocalization-detection system should perform in order to vastly speed up the study of vocal interactions: Detection (whether there is a vocalization), Classification (which type of vocalization it is) and Attribution (which animal vocalized). Automated classification has been done in various species, including rodents (Kobayasi & Riquimaroux, 2012; Soltis, Alligood, Blowers, & Savage, 2012), frogs (Pettitt, Bourne, & Bee, 2012), birds (Giret et al., 2011), bats (Prat, Taub, & Yovel, 2016) and primates (Fuller, 2014; Hedwig, Hammerschmidt, Mundry, Robbins, & Boesch, 2014), including marmosets (Agamaite, Chang, Osmanski, & Wang, 2015; Turesson, Ribeiro, Pereira, Papa, & De Albuquerque, 2016; Zhang, Huang, Gong, Ling, & Hu, 2018). In previous marmoset studies, it was possible to detect the vocalizations by amplitude thresholding of the band- or high-passed audio signal. Attribution was not part of these efforts. As input for classification Agamaite et al extracted 18 acoustic features, chosen by the investigators, in both the time and frequency domain (see Table I in Agamaite et al., 2016). Turesson et al used Linear Predictive Coding filters for feature extraction (Turesson et al., 2016). Zhang et al., (2018) is the most recent work to automatically classify marmoset calls with higher classification accuracy and low frame error rate. They employed a deep recurrent neural network with fully connected layers and LSTM (Long Short Term Memory) (Graves, Mohamed, & Hinton, 2013), in some cases. The data acquisition was from only one animal at a time and thus there is no source attribution or the need to separate background calls. Furthermore, in Zhang et al., the audio recordings were preprocessed using bandpass filters and log-mel filter banks to manually select features to train the network. Call detection and classification were two separate processes and their performance too was evaluated individually.

**Table I.** The 8 call types we distinguish and the ‘noise’ category, containing vocalizations from animals in the background, and sounds that are not from animals. Right column: the number of segments included. (color online)

| Call type | Number of segments |
| --- | --- |
| Trill | 23549 |
| Twitter | 10614 |
| Phee | 10121 |
| Chirp | 5917 |
| Ek | 3265 |
| Trillphee | 2980 |
| Tsik | 2753 |
| Chatter | 1399 |
| Noise (no call type) | 126038 |

The current study is different from previous marmoset studies in several ways. First, recordings were done in a noisy environment. The animals were in their home cage with their cage partner in a room with multiple other cages with animals. While there is benefit in studying animals in conditions where they feel at home and can freely interact with their cage mates, the added noise from animals, cages, air-vents, human personnel, etc. makes it that amplitude thresholding is not sufficient for detecting whether there is a vocalization. Second, our goal was to attribute vocalizations to individuals that are freely moving within the same cage. Attribution is not trivial under these circumstances, because the animals are not spatially separated. We choose to use wearable voice recorders for that reason, and attribute calls using the neural network. The ideal result would be a system where we input two raw audio files and the output is a list of one row per call and columns with start time, stop time, call type, and ID (animal 1 or animal 2).

## III. EXPERIMENT

Our dataset consists of audio recordings done on pairs of animals sharing a cage, each wearing a voice recorder. Between 8 and 20 other animals are present in the room, in other cages. The voice recorders (Polend mini 8GB voice recorder) are mounted at the chest in custom made jackets. The animals are thoroughly habituated to wearing the jackets before recording starts. All animal procedures are overseen by veterinary staff of the MIT and Broad Institute Department of Comparative Medicine, in compliance with the NIH guide for the care and use of laboratory animals and approved by the the MIT and Broad Institute animal care and use committees.

The dataset contains recordings from 16 different individuals in 8 pairs. 36 recording sessions were done, each between 30 and 150 minutes in duration. Total duration of all sessions is 38 hrs. Each recording session yields two mono wav audio files, one for each member of the dyad under study. After each session, the audio files are manually aligned and annotated using Audacity software, version 2.1.3 (Footnote 1). The data are annotated by researchers for the occurrence of 8 different call types: Trill, Twitter, Phee, Chirp, Tsik, Ek, Trillphee and Chatter. Figure 1 shows an example spectrogram of each of the 8 call types.

**Figure 1.**
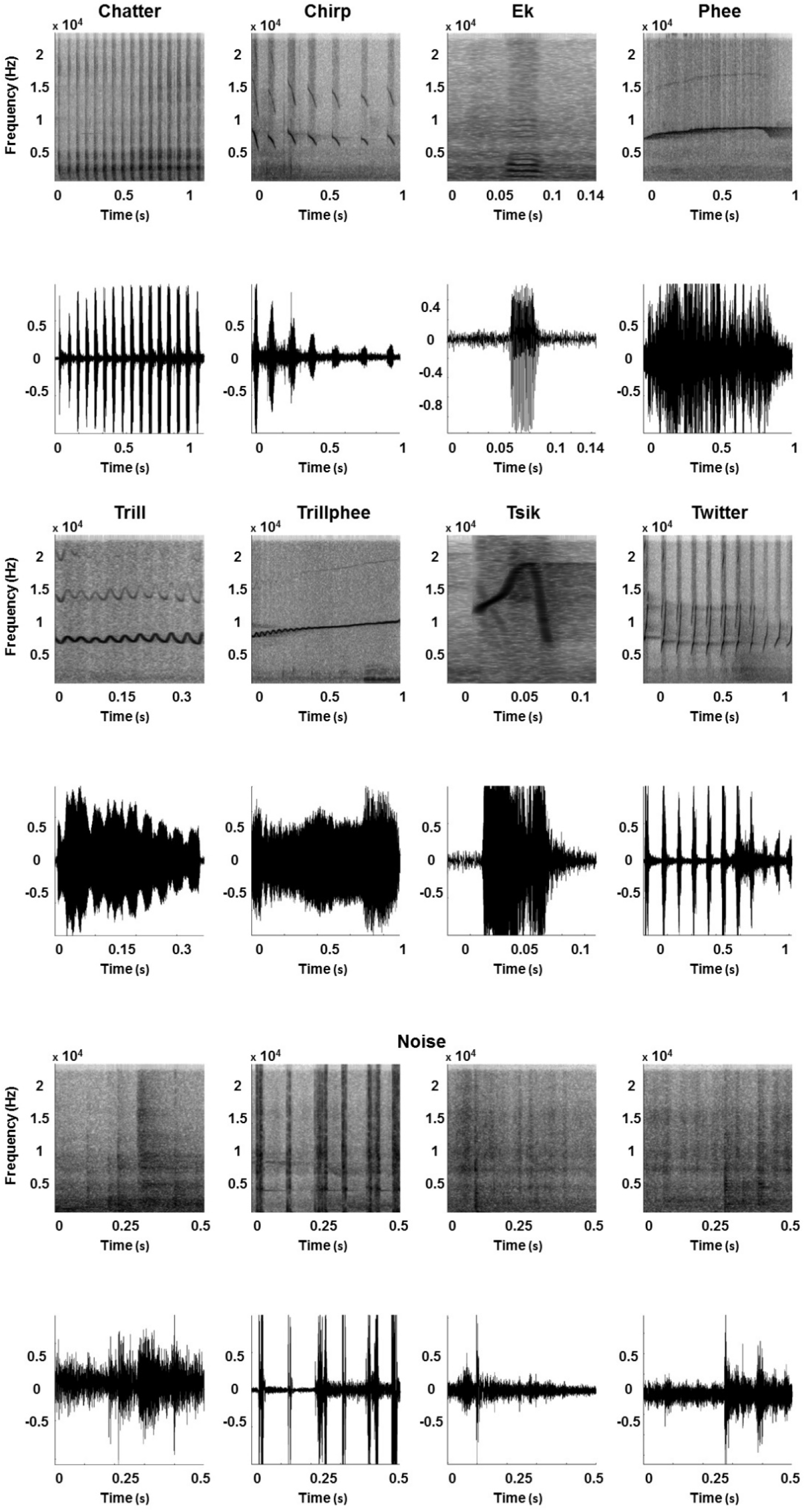
Examples (spectrograms and wave plot) of call types and noises. Row 1-2 (l-r): Chatter, Chirp, Ek, Phee. Row 3-4 (l-r): Trill, Trillphee, Tsik, Twitter. Row 5-6: four different examples of noise.

Annotation is primarily done by inspecting spectrograms. Attribution of calls to either of the two animals wearing a microphone is based on amplitude, reverb, and distinctiveness of the spectrogram image. The calls that can not be attributed are considered to be from other animals in the room and classified as noise. To classify call types we start by using published classifications (Bezerra & Souto, 2008; Epple, 1968; Watson & Buchanan-Smith, n.d.), but some call types are either very uncommon or difficult to distinguish from other call types. For example, we do not distinguish ‘Loud Shrill’, and ‘Seep’ (Watson & Buchanan-Smith, n.d.). For training the network we use the call types for which we have at least 80 exemplars. A total of 15970 marmoset calls labeled by humans are used for training the network.

The audio files are split into 500ms segments with a step size of 150ms. Table 1 shows the number of segments for each call type in the dataset, including the noise category. Spectrograms of the segments are generated using a hamming window, size 512/48000 s and a step size of 92/48000 s, which yields spectrogram images of size 257×256. To each pair of spectrograms, one for each simultaneously recorded channel, we assign a label based on whether there is a human annotation of a call in the middle 150ms of the window, including the call type and which animal does the call (ch 1 or ch 2). If no labeled call is present in the segment, the pair is labeled as noise. Since the vocalizations are brief and there is some time in between vocalizations, the data is heavily unbalanced with vast majority of the labels being noise. To alleviate this, we only include 20% of the segments labeled as noise into the dataset. We use 3 sessions for evaluation and another 3 full sessions for testing. The training dataset is composed of the remaining 30 sessions.

The model is an end to end feedforward convolutional neural network with 4 blocks of 2 convolutional layers followed by a max pooling layer for each of the two spectrogram images that are fed in simultaneously. The convolutional streams are processed separately, after which the two streams are concatenated and followed by two fully connected layers. Each layer uses rectified linear units (Nair & E. Hinton, 2010) as activation functions, except for the final layer which uses softmax function. Figure 2 shows the architecture of our standard model. Each model we test shares this architecture except for when stated otherwise.

Our models are trained for 74000 batches of 25 examples each using Adam optimizer (Kingma & Ba, 2015) and a cross entropy loss function with a learning rate of 0.0003 exponentially decreased by multiplying with 0.97 after every 2000 batches and epsilon of 0.001. Figure 3 shows training and evaluation accuracy during training of the standard model.

**Figure 2.**
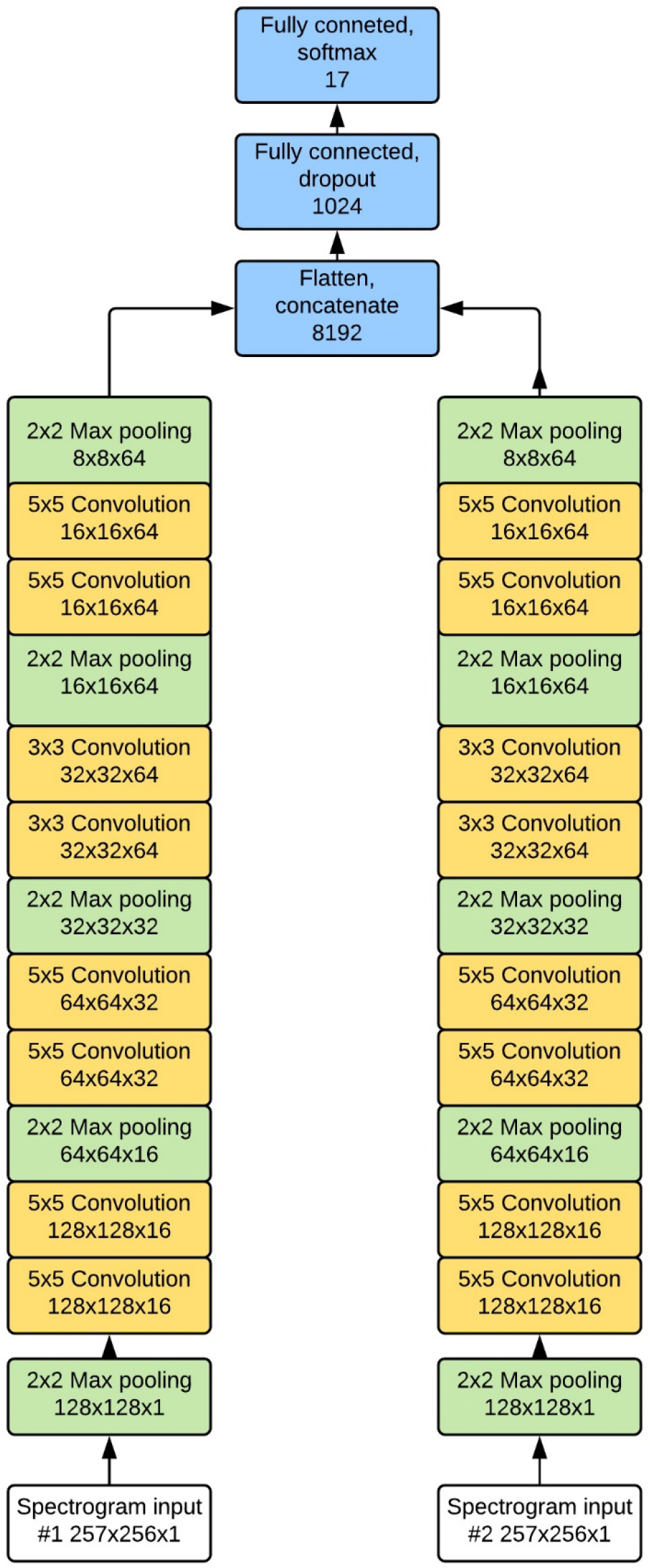
The architecture of the standard dual stream version of our network. The kernel size of each layer is shown before the type of the layer and the dimensions of the layer(excluding batch) are below. Strides of 1 are used for each convolutional layer and strides of 2 for each max pooling layer. (color online)

**Figure 3.**
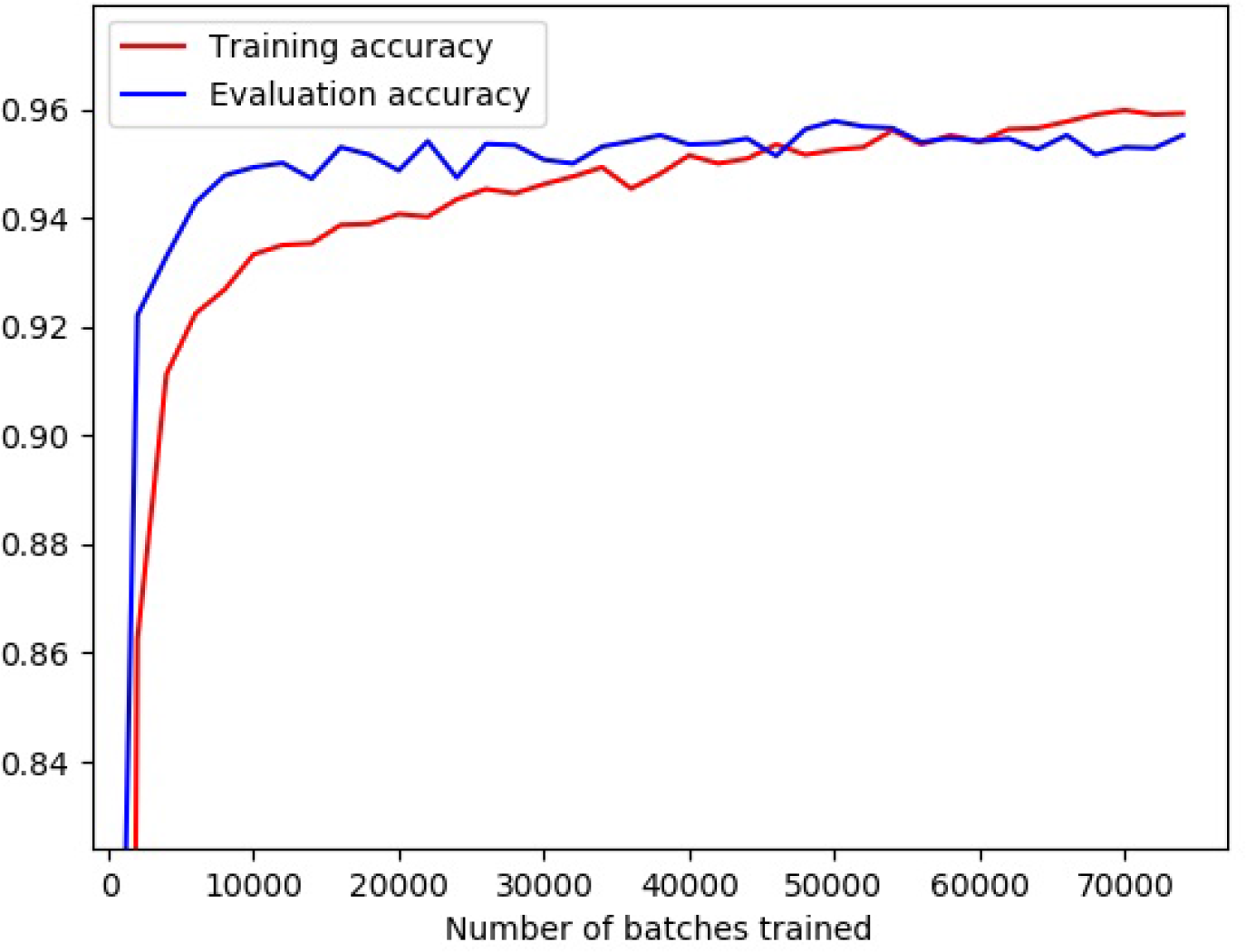
Development of training and evaluation accuracies of the standard network (color online)

Since our network does classification and attribution at the same time, the design of the final layer can be tailored accordingly. We experiment with three different layouts for the final layer, shown in Figure 4. The final layers with 17 and 9+3 units can only a detect a call from one animal at a time, so we also test a multilabel version of the 17 unit final layer that uses sigmoid activation functions for the final layer instead of softmax that can detect several calls at a time. For predicting with the multilabel network we only look at the highest prediction of the network for each animal since it is not possible for a single animal to make two calls at the same time. We also test how beneficial it is to use two input streams by comparing our results against those achieved by training a network of the same structure but with only one input stream being fed one of the audio files at a time and only classifying the call made by that animal.

**Figure 4.**
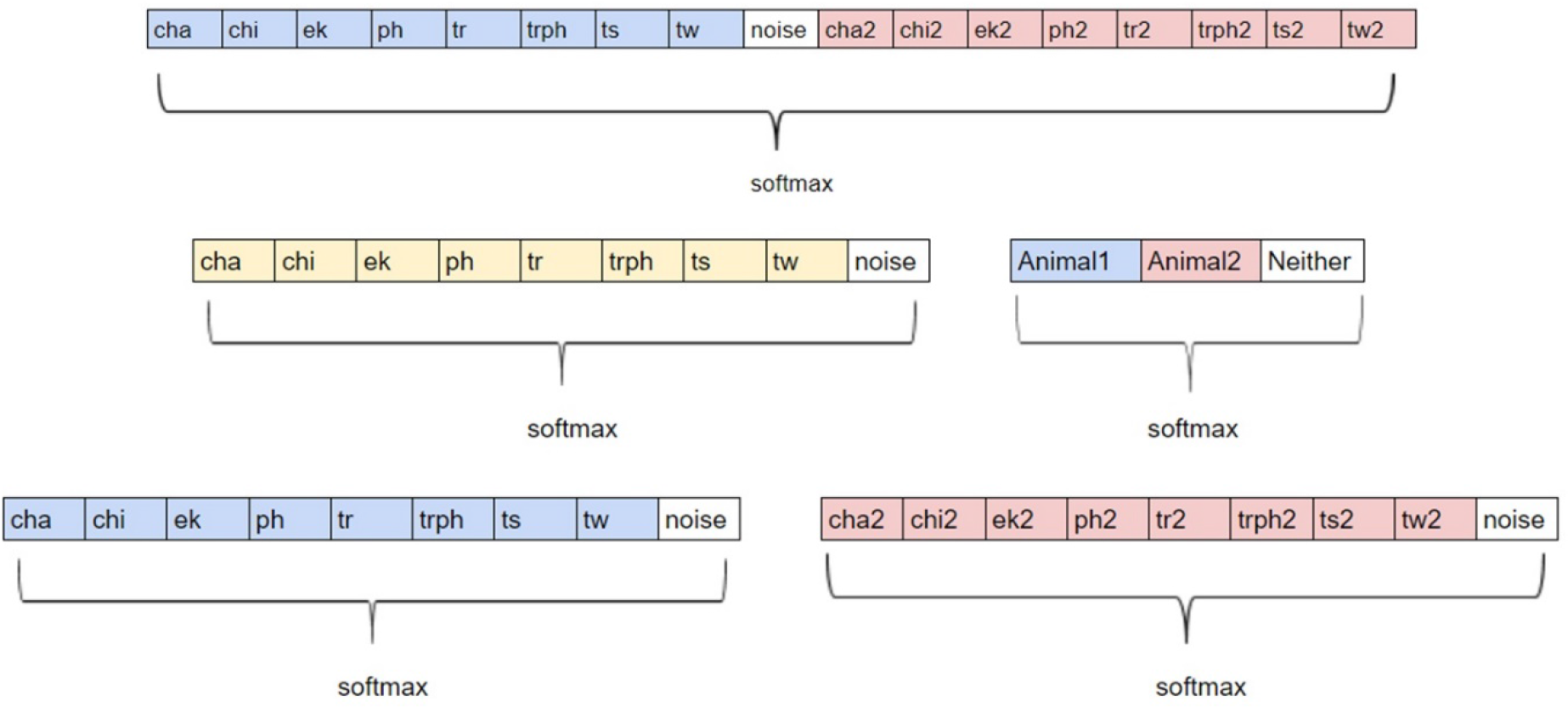
Top: The standard 17 unit final layer. Middle: The 9+3 final layer setup. Bottom: 9+9 final layer setup, capable of detecting two calls at the same time unlike the other final layer options. (color online)

We explore various methods of data augmentation including translations, adding random noise and multiplying the input by a random value close to one. However, none significantly improve our results. We use a small random vertical and horizontal roll (shifting the array while feeding overflowing values on the other end) of up to five pixels each since that improved results on a smaller dataset. The effects are small on the actual dataset. Besides that, we randomize which input gets fed into input1 and input2 and adjust the labels accordingly, in order to try and keep the network “speaker” independent instead of learning to recognize each monkey or recording. The standard version of our network does not use batch normalization (Sergey Ioffe & Christian Szegedy, 2015) but we also test models using batch normalization after every weighted layer before applying the nonlinearity.

The goal of our project is to detect, classify and attribute each call in a pair of long recordings, which includes getting a good temporal accuracy. To evaluate performance we run the network with 500ms window and 50ms step size on the test data. To produce each prediction we take the average of the predictions using a window centered around the 50ms we are predicting and the windows shifted by 50ms and 100ms to both directions. We then apply a cutoff to this average, such that predictions with confidence less than a certain cutoff were classified as noise. This is done for all call types except trillphee, which is a mix between trill and phee: If the highest prediction is trill or phee, we combine the confidence of that and trillphee before applying the cutoff, and if highest is trillphee, we sum it together with trill and phee before applying cutoff. We found that cutoff values around 0.8 produce the best results.

The network’s performance is measured by discretizing the ground truth into 50ms segments and labeling each of these according to the original labels. We then test the accuracy of our model on this data. Table II shows the definitions of the metrics we use to evaluate our results. F1-score is the main metric we use when comparing performance of different models.

**Table II.** Definition of terms

| Term | Explanation |
| --- | --- |
| True positive(TP) | There is a call in the frame and it was correctly classified by the network |
| False positive(FP) | There is no call in the frame but the network predicted a call |
| True Negative(TN) | There is no call and the network predicted no call(=noise) |
| False Negative(FN) | There is a call but the network predicted a wrong call, or no call |
| Recall | $\#TP/(\#TP+\#FN)$ |
| Precision | $\#TP/(\#TP+\#FP)$ |
| F1-score | $2*Recall*Precision/(Recall+Precision)$ |
| Accuracy | Standard framewise accuracy, $(\#TP+\#TN)/(\#TP+\#FP+\#TN+\#FN)$ |

## 4. RESULTS

We test different versions of our network using cutoff values of 0,0.3,0.4,0.5 and 0.6-0.95 with increments of 0.05. Figure 5 shows how changing the cutoff affects recall, precision and F1-score.

**Figure 5.**
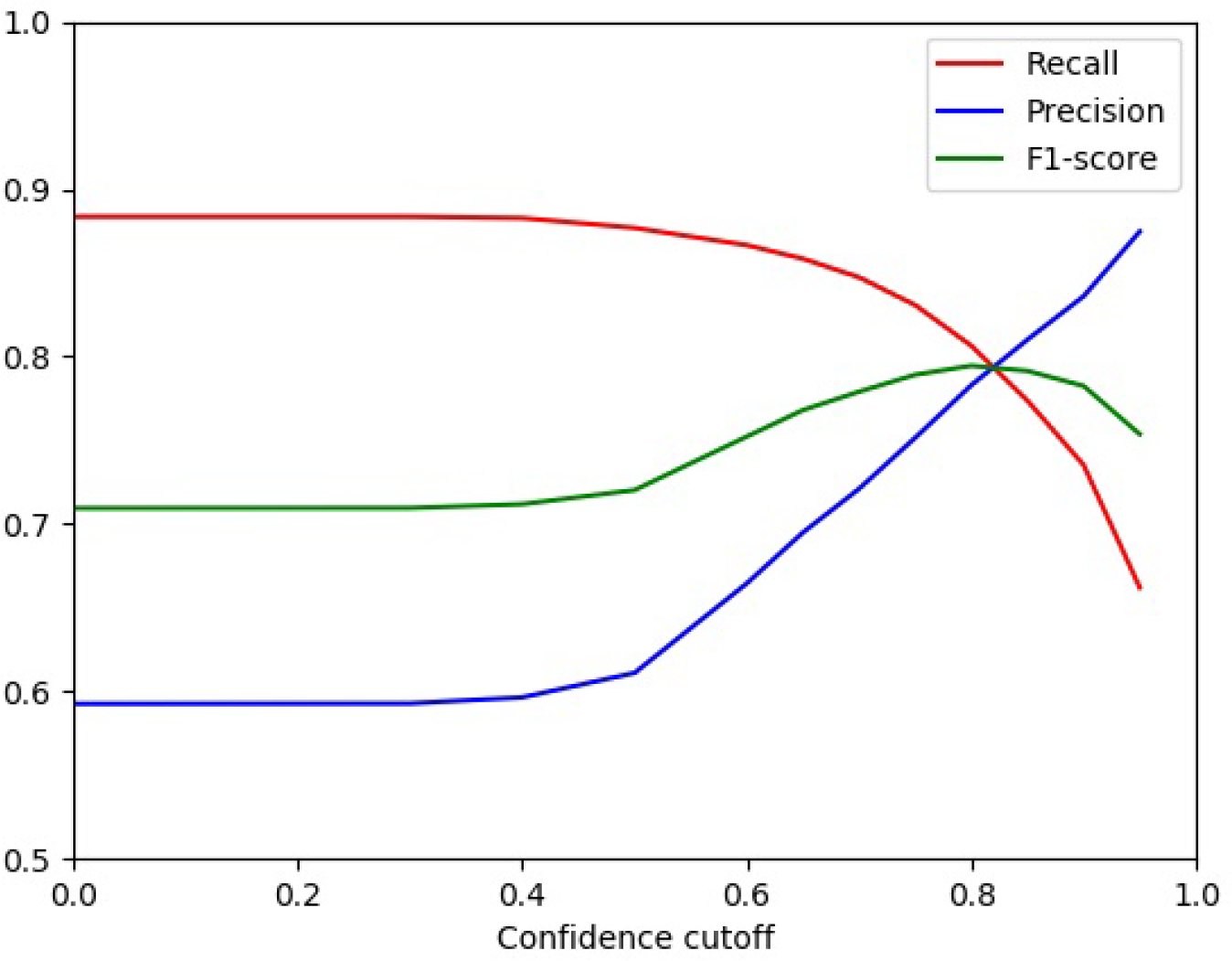
Recall, precision, and F1-score of the standard model as a function of cutoff. (color online)

Table III shows the best F1-score obtained by each version among the tested cutoffs. As a baseline example we also train a single stream model with the AlexNet (Krizhevsky, Sutskever, & Hinton, 2012) architecture on our data.

**Table III.**
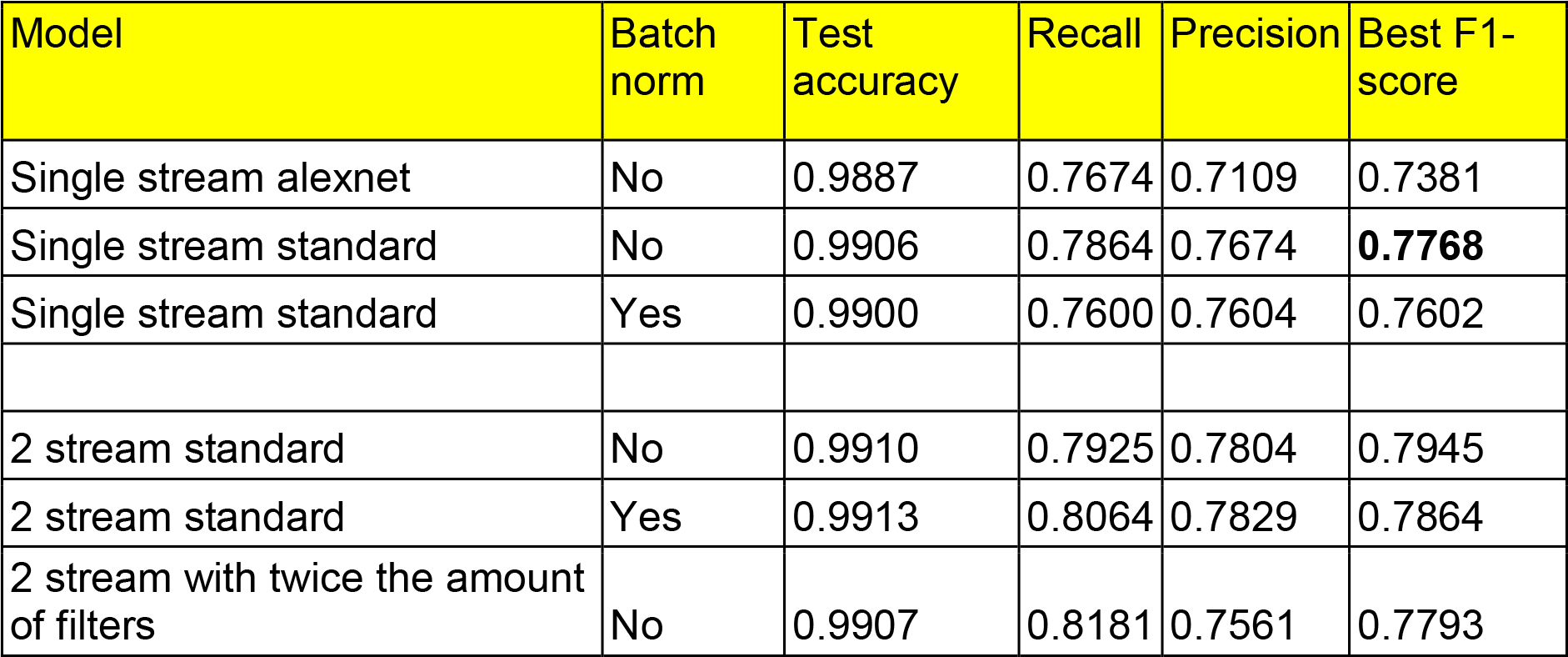
Batch normalization, Test accuracy, Recall, Precision and Best F1-scores for all models tested (color online)

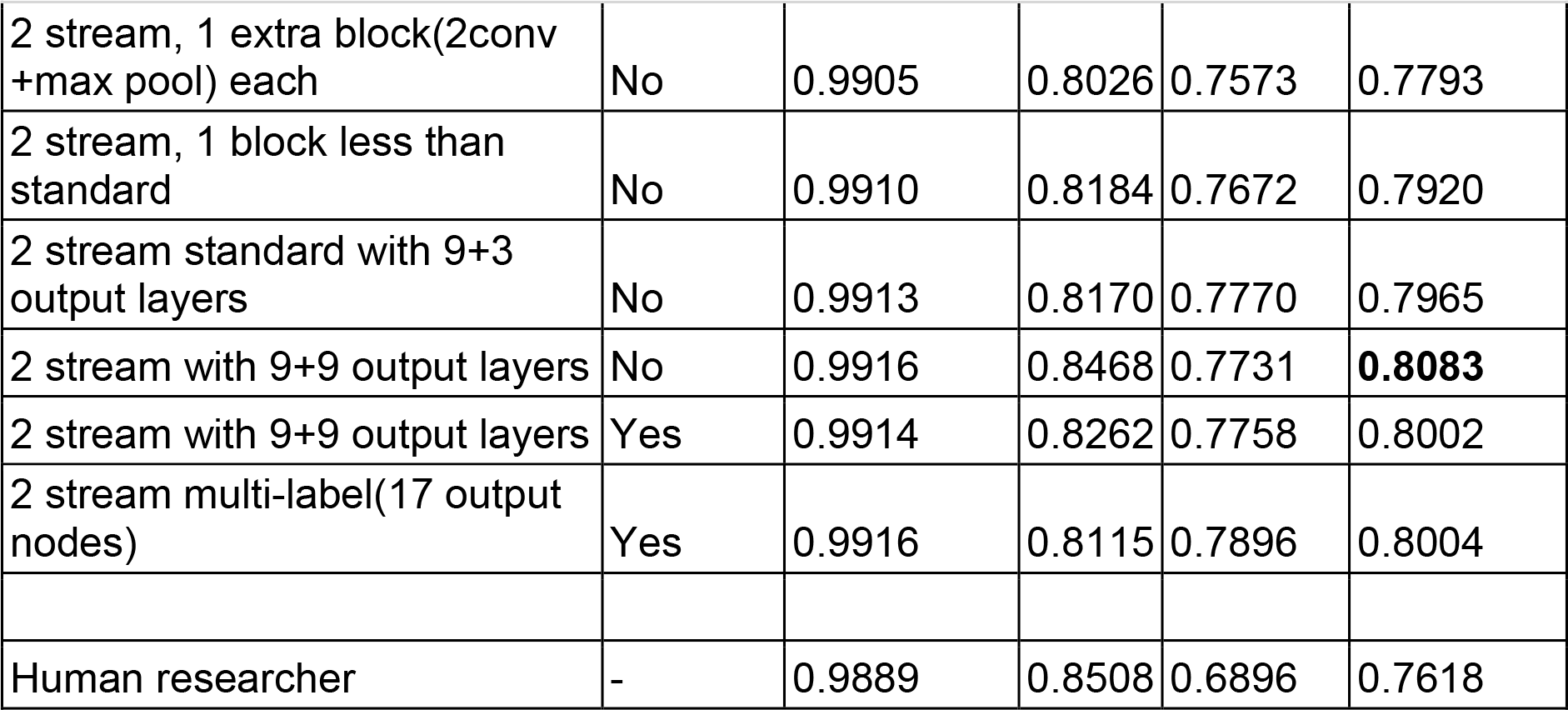

Our best performing model is able to achieve an F1-score of 0.8083, and a framewise accuracy of 0.9916 on the test set, with the accuracy being significantly higher than F1-score due to vast majority of frames being noise. We can see that small changes to network architecture do not result in very significant differences in final performance, all 9 networks utilizing dual audio input reach F1-scores between 0.7793 and 0.8083. Interestingly, batch normalization (‘batch norm’ column in Table III) does not improve the performance of our network and instead seems to result in slightly lower accuracies. The layout of the final layer does not seem to noticeably affect network performance, with the exception that networks capable of detecting two simultaneous calls seem to perform slightly better than ones that can not, with all of those reaching F1-scores over 0.8 while none of the ones that can only detect a single call at a time do. Figure 6 shows an example of the network detecting two different calls at the same time.

**Figure 6.**
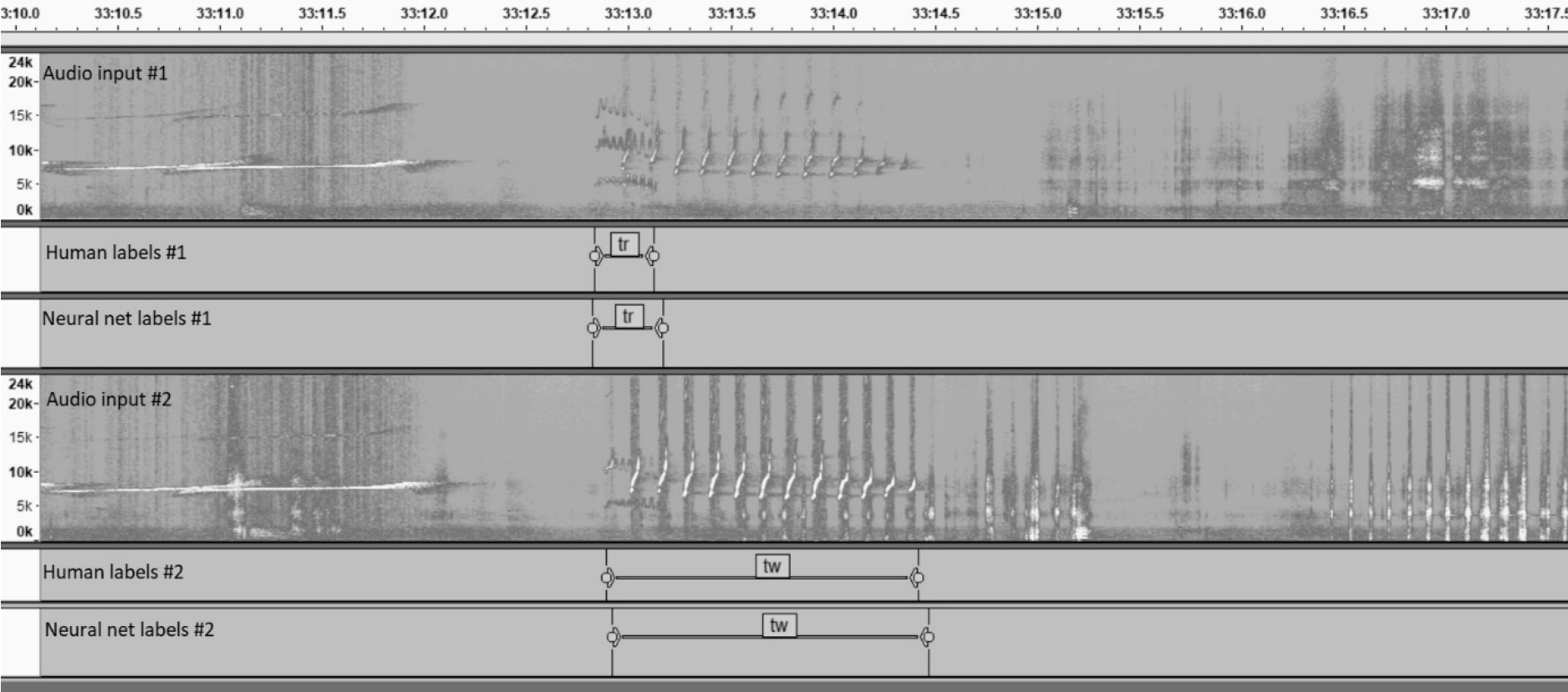
An example of our best performing model detecting two different calls simultaneously on a test session. The recording is from two marmosets housed in the same cage, each wearing a microphone located at the chest (as all the recordings in our dataset). The top spectrogram shows animal 1, and the bottom spectrogram shows animal 2. Many marmoset calls are visible in both spectrograms, yet certain marmoset calls are clearly more pronounced on one channel than on the other. Between time points 33.10 and 33.12 there is a ‘phee’ call which is equally pronounced on both mics, and likely was produced by an animal in another cage. At 33.13, a ‘trill’ call occurs. It is most pronounced on ch1, and this was labeled as a call coming from animal 1, but both human observer and the network. At the same time, a ‘twitter’ call begins. This is more pronounced on ch2, and is labeled as such by both human and network. This figure was created using Audacity software (footnote 1)

Our results show that using two audio streams improves performance. Each network that uses two input channels beats every network that uses only one input. The margin is not very wide, the best performing single stream reaches an F1-score only 0.0025 lower than the worst performing two stream network’s score. This difference is most likely smaller because the networks with single input can detect two simultaneous calls while the worst performing two stream networks can not. Regardless, the difference is clear enough to see that two inputs are beneficial.

Our network is trained to replicate the labels of human observers, yet between human observers, there is inherent variability. Our database is annotated by only a single observer per observation. To better appraise the performance of the network, it is useful to know whether the difference between the network and the human observer is bigger or smaller than the difference between human observers. Therefore, we ask a different human observer to re-label the test sessions and test how closely these labels match the original observer using the same metrics we used for measuring the performance of our network. The result shows that the second human observer was less accurate than our network in replicating the original labels (‘human researcher’ Table III). This underscores that there is variability between human observers, and that our network can perform better than some human observers given this set of training data. Inspired by these results we also measure accuracy again using the labels created by the second human observer (human relabel) as ground truth instead of the original ones. The results of this experiment are shown in table IV. Our best network achieves an F1-score of 0.7858 when compared to the human relabels, while the original human labels only reaches an F1-score of 0.6694 in this comparison. This shows that our network is closer to both human labels than they are to each other, indicating it can capture features common between human labels, with less variance, suggesting that its results are better than the average human, in effect surpassing the accuracy of its training set labels.

**Table IV.** F1-scores for human relabels and our best model.

|  |  |  |  |  |
| --- | --- | --- | --- | --- |
|  |  |  | **Predictions:** |  |
|  |  | Original human labels | Human relabels | Our best model |
|  | Original human |  | F1-score | F1-score |
|  | labels | - | 0.7618 | 0.8083 |
| **Ground truth:** |  |  |  |  |
|  | Human relabels | F1-score |  | F1-score |
|  |  | 0.6694 | - | 0.7858 |

We also test our network on the marmoset call dataset shared introduced in Turesson et al. (2016). This dataset consists of 321 segmented calls, so it is much smaller than our dataset, and because the calls are pre-segmented, the task is one of classification, not detection. Our network is optimized for larger datasets and as such has the risk of overfitting to smaller dataset, but we are able to achieve good results. We train a single stream normalized version of our network from scratch for 600 batches of 25 examples 10 times while using randomly selected 90% of the calls in each call type for training and evaluated accuracy on the remaining 10%. For training we create spectrogram images of 500ms sliding window with 200ms steps over the duration of each call and label each image with the label of the call. For prediction we run the network with each window of the call and use the mean of those as our prediction. Table V shows our results measured by the metrics defined by Turesson et al. We find out that our network’s accuracy is better than the best model tested by Turesson et al., which is an SVM model with Linear Predictive Coding.

**Table V.** Performance of our network on Turesson et al’s dataset and comparison with their model. (color online)

| Model | F1-score | Accuracy |
| --- | --- | --- |
| Our network trained from scratch | 0.890 | 0.936 |
| SVM with LPC(their best) | 0.852 | 0.843 |

We implement our network into an easy to use program that can classify marmoset recordings. We also optimize its performance which allows our network to classify an hour long audio recording in less than 15 minutes using a high end laptop. Our model as well as source code used for the results in this paper and a subset of our data are freely available at https://marmosetbehavior.mit.edu/

## V. DISCUSSION

The end-to-end feedforward CNN introduced in this article is one of a kind, in that it is capable of automatic call detection, classification of call types, and attribution of the caller, all together. The network automatically detects whether any given segment of hour long audio files of dual channel marmoset vocal recordings contains a call or not. It then classifies the detected calls into one of 9 types (8 call types + 1 noise category), and finally attributes the identity of the call to the animals wearing the microphone (either of the two animals or neither). In addition, we have adapted the CNN for single stream audio recordings as well. To our knowledge, this task optimized feedforward CNN gives the best performance to date on automatic vocal classification of marmoset calls.

Unlike previous efforts that classify call types on noise-free, pre-segmented audio (Agamaite et al., 2015; Turesson et al., 2016), our network uses raw, noisy spectrograms. The network is thus robust in detecting and classifying in an environment that is noisy, and doesn’t require any preprocessing of the input audio stream. Agamaite et al. (2015) describe a quantification of audio features, chosen by the investigators, to identify different call types. Here we let the network learn the useful features that it needs to enable the trio of detection, classification, and caller attribution.

Our approach is image-based, using the spectrogram of the audio. However, our dataset is different from many other image datasets. Typically, large benchmark datasets like ImageNet (with 15 million images, and 1000s of categories) used in testing and validating machine learning algorithms and neural networks are well curated. New object recognition algorithms typically use a uniform number of each of the different categories to train their networks. Unlike such datasets, our training set is quite modest in size and is not curated (noise-free, anti-aliased, mean-centered etc.,), and not trained with a uniform number of samples of different calls (eg., more ‘trill’, or ‘phee’, than ‘chatter’), because not all calls are equally common, yet all calls are potentially important to study.

Turesson et al. (2017) used a very small, but well curated dataset (321 calls) which they have made freely available. They applied several machine learning and neural network algorithms and found that an SVM (Support Vector Machine) with Linear Predictive Coding (LPC) had the best performance metrics. Training our network from scratch achieved results near to their best performance. However, when we did transfer learning of our network to their small training dataset, the performance was much higher than what they had reported with their best methods. This shows that our convolutional network contains robust learned representations and that it can classify marmoset calls recorded in different environmental scenarios. We believe even better results could be achieved with this dataset using our model after careful hyperparameter optimization. Besides other marmoset datasets, it is very well possible that the convolutional network we have designed here is capable of transfer learning to be adapted for vocal classification in other species. Our shared code is freely available for others to use.

Our study diverges from Zhang et al. (2018) in that we recorded from two animals instead of one which is a step towards analyses on social behavior, but required us to solve the problem of source attribution. Similar to Agamaite et al., (2015), Zhang et al., hand-picked three short-time acoustic features that are extracted for each audio frame: energy, peak-ratio of autocorrelation (PRA), and log mel-filter bank spectrum. It is these features that are used for detecting voiced vs non-voiced segments, and later to train the network on the extracted log mel-filter bank spectra. The call detection employed was a rule-based threshold detection algoithm that is applied in many human speech processing systems for voice activity detection (VAD). Thus call detection and classification were done separately, with a rule-based approach for detection followed by a neural network training for call classification. In our approach, we directly feed our network raw, noisy spectrograms from the dual channels that contain background animal calls, and a variety of noise in the environment. Detection, classification, and attribution are performed by a single network and we avoid the possible bias that could be introduced by hand-picking features. The neural network of Zhang et al., is a fully connected recurrent neural network (RNN) with LSTM, while ours is a feedforward deep convolutional neural network. We found that adding LSTM layers to our system does not improve the performance on the task.

Many systems have applied the architecture of AlexNet (Krizhevsky et al., 2012) to a high degree of success. In that vein, we trained an Alexnet model on a single stream of our dataset with the raw spectrograms as the input images. The performance of this network was much worse than our convolutional network (both single and dual stream audio). This result and the transfer learning of our network for the data set from Turesson et al. (2017) shows that as a proof of principle, we have a task optimized convolutional network that has learned features generalized over the space of vocal calls.

We show that our network more closely replicates labelling from a human observer than a second human observer is able to replicate the first human observer. This highlights that there is considerable variation in human labelling, even among experts. In the existing literature there is considerable variation in the definition and number of call types distinguished (Bezerra & Souto, 2008; Epple, 1968)(Epple, Bezerra, watson, Agamaite, Turesson).

Further improvement of auto-detection and classification efforts will be aided by standardization of the definition of call types (eg., ‘phee’, ‘short phee’, ‘long phee’ etc.,), naming conventions, and a robust, yet flexible dataset (with noise, multiple streams etc.,). Unsupervised clustering of audio might be worth exploring as a way to establish objective and bias-free categorization. Additional areas of improvements include having an expanded training set, labels, timings of the calls corroborated by multiple humans to increase reliability.

Footnote 1: Audacity® software is copyright © 1999-2018 Audacity Team. The name Audacity® is a registered trademark of Dominic Mazzoni.

## REFERENCES

Agamaite, J. A., Chang, C.-J., Osmanski, M. S., & Wang, X. (2015). A quantitative acoustic analysis of the vocal repertoire of the common marmoset (Callithrix jacchus). The Journal of the Acoustical Society of America, 138(5), 2906–2928. https://doi.org/10.1121/1.4934268

Bezerra, B. M., & Souto, A. (2008). Structure and Usage of the Vocal Repertoire of Callithrix jacchus. International Journal of Primatology, 29(3), 671–701. https://doi.org/10.1007/s10764-008-9250-0

Boddapati, V., Petef, A., Rasmusson, J., & Lundberg, L. (2017). Classifying environmental sounds using image recognition networks. In Procedia Computer Science. https://doi.org/10.1016/j.procs.2017.08.250

Eliades, S. J., & Miller, C. T. (2017). Marmoset vocal communication: Behavior and neurobiology. Developmental Neurobiology. https://doi.org/10.1002/dneu.22464

Epple, G. (1968). Comparative studies on vocalization in marmoset monkeys (Hapalidae). Folia Primatologica; International Journal of Primatology, 8(1), 1–40. Retrieved from http://www.ncbi.nlm.nih.gov/pubmed/4966050

Fuller, J. L. (2014). The vocal repertoire of adult male blue monkeys (Cercopithecus mitis stulmanni): A quantitative analysis of acoustic structure. American Journal of Primatology, 76(3), 203–216. https://doi.org/10.1002/ajp.22223

Giret, N., Roy, P., Albert, A., Pachet, F., Kreutzer, M., & Bovet, D. (2011). Finding good acoustic features for parrot vocalizations: The feature generation approach. The Journal of the Acoustical Society of America, 129(2), 1089–1099. https://doi.org/10.1121/1.3531953

Graves, a, Mohamed, A., & Hinton, G. (2013). Speech recognition with deep recurrent neural networks. Icassp. https://doi.org/10.1109/ICASSP.2013.6638947

Hedwig, D., Hammerschmidt, K., Mundry, R., Robbins, M. M., & Boesch, C. (2014). Acoustic structure and variation in mountain and western gorilla close calls: a syntactic approach. Behaviour, 151, 1091–1120. https://doi.org/10.1163/1568539X-00003175

Henry, L., Craig, A. J. F. K., Lemasson, A., & Hausberger, M. (2015). Social coordination in animal vocal interactions. Is there any evidence of turn-taking? The starling as an animal model. Frontiers in Psychology, 6, 1416. https://doi.org/10.3389/fpsyg.2015.01416

Jennings, C. G., Landman, R., Zhou, Y., Sharma, J., Hyman, J., Movshon, J. A., … Feng, G. (2016). Opportunities and challenges in modeling human brain disorders in transgenic primates. Nature Neuroscience, 19(9), 1123–1130. https://doi.org/10.1038/nn.4362

Kingma, D. P., & Ba, J. L. (2015). Adam: a Method for Stochastic Optimization. International Conference on Learning Representations 2015. https://doi.org/http://doi.acm.org.ezproxy.lib.ucf.edu/10.1145/1830483.1830503

Kobayasi, K. I., & Riquimaroux, H. (2012). Classification of vocalizations in the Mongolian gerbil, Meriones unguiculatus. The Journal of the Acoustical Society of America, 131(2), 1622–1631. https://doi.org/10.1121/1.3672693

Krizhevsky, A., Sutskever, I., & Hinton, G. E. (2012). ImageNet Classification with Deep Convolutional Neural Networks. Advances In Neural Information Processing Systems. https://doi.org/http://dx.doi.org/10.1016/j.protcy.2014.09.007

Levinson, S. C., & Torreira, F. (2015). Timing in turn-taking and its implications for processing models of language. Frontiers in Psychology, 6, 731. https://doi.org/10.3389/fpsyg.2015.00731

Miller, C. T., Freiwald, W. A., Leopold, D. A., Mitchell, J. F., Silva, A. C., & Wang, X. (2016). Marmosets: A Neuroscientific Model of Human Social Behavior. Neuron, 90(2), 219–233. https://doi.org/10.1016/j.neuron.2016.03.018

Miller, C. T., Mandel, K., & Wang, X. (2010). The communicative content of the common marmoset phee call during antiphonal calling. American Journal of Primatology, 72(11), 974–980. https://doi.org/10.1002/ajp.20854

Nair, V., & E. Hinton, G. (2010). Rectified Linear Units Improve Restricted Boltzmann Machines Vinod Nair. Proceedings of ICML (Vol. 27).

Pettitt, B. A., Bourne, G. R., & Bee, M. A. (2012). Quantitative acoustic analysis of the vocal repertoire of the golden rocket frog (Anomaloglossus beebei). The Journal of the Acoustical Society of America, 131(6), 4811–4820. https://doi.org/10.1121/1.4714769

Prat, Y., Taub, M., & Yovel, Y. (2016). Everyday bat vocalizations contain information about emitter, addressee, context, and behavior. Scientific Reports, 6(1), 39419. https://doi.org/10.1038/srep39419

Sacks, H., Schegloff, E. A., Jefferson, G., & SacksEmanuel SchegloffGail Jefferson, H. A. (1974). A simplest systematics for the organization of turn-taking for conversation, 50(21), 696–73550. https://doi.org/10.1353/lan.1974.0010

Sergey Ioffe, G., & Christian Szegedy, G. (2015). Batch Normalization. Icml. https://doi.org/10.1007/s13398-014-0173-7.2

Soltis, J., Alligood, C. A., Blowers, T. E., & Savage, A. (2012). The vocal repertoire of the Key Largo woodrat (Neotoma floridana smalli). The Journal of the Acoustical Society of America, 132(5), 3550–3558. https://doi.org/10.1121/1.4757097

Turesson, H. K., Ribeiro, S., Pereira, D. R., Papa, J. P., & De Albuquerque, V. H. C. (2016). Machine learning algorithms for automatic classification of marmoset vocalizations. PLoS ONE. https://doi.org/10.1371/journal.pone.0163041

Watson, C. F. I., & Buchanan-Smith, H. M. (n.d.). MarmosetCare.com. Retrieved January 1, 2018, from http://www.marmosetcare.com/

Zhang, Y.-J., Huang, J.-F., Gong, N., Ling, Z.-H., & Hu, Y. (2018). Automatic detection and classification of marmoset vocalizations using deep and recurrent neural networks. The Journal of the Acoustical Society of America, 144(1), 478–487. https://doi.org/10.1121/1.5047743

